# PackIO and EphysViewer: software tools for acquisition and analysis of neuroscience data

**DOI:** 10.1101/054080

**Authors:** Brendon O. Watson, Rafael Yuste, Adam M. Packer

## Abstract

We present an open-source synchronization software package, PackIO, that can record and generate voltage signals to enable complex experimental paradigms across multiple devices. This general purpose package is built on National Instruments data acquisition and generation hardware and has temporal precision up to the limit of the hardware. PackIO acts as a flexibly programmable master clock that can record experimental data (e.g. voltage traces), timing data (e.g. event times such as imaging frame times) while generating stimuli (e.g. voltage waveforms, voltage triggers to drive other devices, etc.). PackIO is particularly useful to record from and synchronize multiple devices, for example when simultaneously acquiring electrophysiology while generating and recording imaging timing data. Experimental control is easily enabled by an intuitive graphical user interface. We also release an open-source data visualisation and analysis tool, EphysViewer, written in MATLAB, as well as a module to import data into Python. These flexible and programmable tools allow experimenters to configure and set up customised input and output protocols in a synchronized fashion for controlling, recording, and analysing experiments.

## Introduction

Experimental protocols in neuroscience often require recording data simultaneously from many different sources, such as electrophysiological, behavioral, and synchronization data from various pieces of equipment. Additionally, data often needs to be generated to drive or trigger equipment such as electrophysiological amplifiers, imaging systems, and stimulus delivery equipment. There are several commercial and open-source packages that address these needs but each has limitations such as their lack of flexibility, high cost or the limited repertoir of protocols it can perform.

We present here an open-source LabVIEW software package (“PackIO”) for acquiring and generating data from National Instruments cards. It is meant to trigger and synchronize equipment as well as record any data, with a few specific modules for electrophysiology. All protocols can be driven in a hardware-timed fashion, limited only by the specifications of the chosen National Instruments data acquisition card. This software has already been used in 32 publications^1^^−^^32^ performing both *in vitro* and *in vivo* experiments from several different labs. We also present additional open-source software, EphysViewer, which was written in MATLAB and provides a suite of analysis tools for detailed investigation of data acquired with PackIO. In combination, these software packages enable experiments requiring acquisition and generation of data and can be triggered in several modes.

## Results

### Concept

PackIO is designed to perform simultaneous equipment control and data recording in a modular fashion. We provide a graphical user interface designed for a variety of experimental types with specialized tools for electrophysiology. As a result of the modular design, we provide generic but customizeable control options allowing users to specify timing and amplitudes of outputs while also allowing selection of recording inputs. Outputs can be sequenced and a Producer/Consumer software design (below) allows multiple simultaneous loops and timing structures to interact during an ongoing experiment. Relatedly, experiments can be run either continuously or in tracked and timed episodes. All experimental inputs and outputs are visualized in real time and can be stopped and revised as necessary with intuitive graphical tools. Finally, a MATLAB visualization tool is provided, EphysViewer, which is capable of displaying any vectorized or timeseries data, but which is also customized to vizualize experiments executed in PackIO. Given the flexible and intentionally generic nature of this tool, we will specify below use parameters and concepts for users.

### Outline of PackIO workflow

When PackIO starts, the main window appears, which is where experiments will be started, monitored online, and stopped. Upon first opening, the Master Setup window also appears in order to set up an experimental protocol.

The workflow is as follows:

1. Set up an experiment in the Master Setup window.
2. Click start in the main window to begin acquisition and/or generation of data.
3. Click stop to conclude the experiment.
4. Optionally repeat steps 2-3 to run the same experiment again.
5. Click setup in the main window to restart at step 1 if desired, or click close to exit the program.

### Running PackIO

When first run, PackIO will create directories in the directory the main program library is in (C:\PackIO is recommended if using the Windows operating system) if they do not already exist. These directories are ‘TempData’ (temporary data files for the ‘Go back’ feature) and ‘Configurations’ which contains four subdirectories: Epoch, Input, Output and Seal, which contain configuration files for Epoch Loader, Master Setup input tab, Master Setup output tab, and Seal Test (respectively).

### PackIO Main window

The main window of PackIO (**Fig. 1**) is where all the major features of the program can be accessed. On the left side are seven numbered main displays which show incoming data. The controls on the right are described further below.

**Figure 1:**
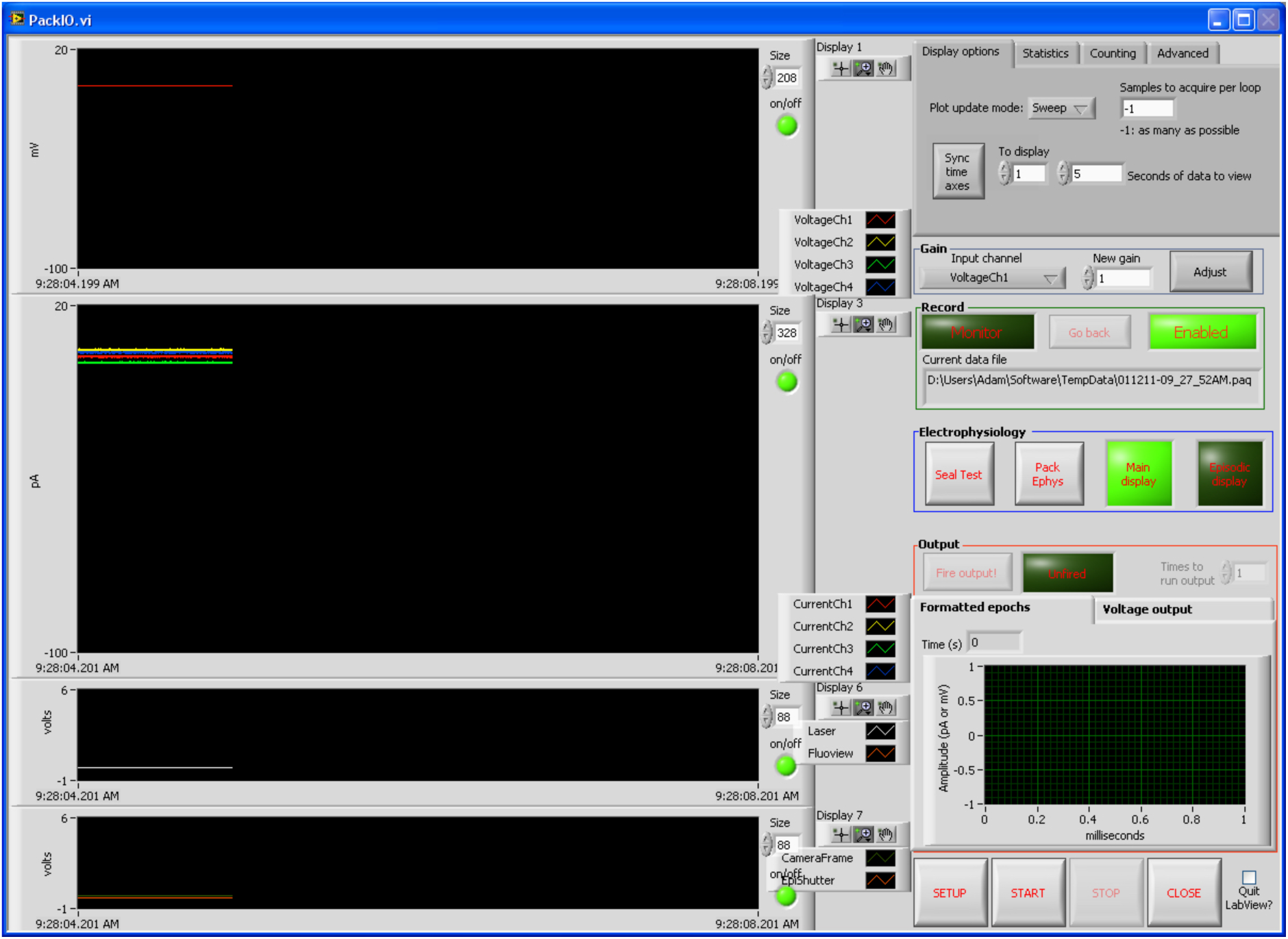
Main window of PackIO. Experiments are started and stopped here. The setup, seal test, PackEphys, and episodic modules are accessed here as well. Incoming data is plotted based on the selections made in the master setup window.

Display options: ‘Plot update mode’ controls the manner in which data is updated to the main displays at the left. ‘Samples to acquire per loop’ is the number of samples the to read from every channel of an acquisition during each loop of the main program’s operation. The default of −1 will acquire as many samples as are available. ‘Sync time axes’ locks the x-axes of the main display windows to the x-axis shown in the ‘To display’ box. Seconds of data to view indicates how much time to display in the main displays.

Statistics: displays information regarding the condition of PackIO, which is useful when diagnosing issues with slower computers.

Counting: implements DAQ counter functions. The count is displayed and can be divided by a user-set value.

Advanced: allows the user to set a value to which an output channel will be set after completion of a task. Normally, this value is zero, but there are instances, for example when controlling certain pieces of equipment such as laser controllers, in which a negative value may be desired.

Gain controls: The gain of an input channel can be adjusted during a recording to the value in ‘New gain’ by clicking the Adjust button.

Recording controls: If the button on the upper left is not lit, it will read ‘Monitor’ which indicates no data will be recorded to a file selected by the user (but data may still be recorded if the ‘Go back’ feature is enabled, see below.) If the ‘Monitor’ button is clicked, it will become lit and read ‘Record’ in which case the user will be prompted for a file name when START is clicked and recording commences. It is recommended to use the file extension *.paq when recording PackIO data. The ‘Go back’ feature records all data, even when the user does not explicitly specify to record. The user can then click ‘Go Back’ if the user decides he or she actually wants to save the data just displayed. The file name of the data that has been stored is then displayed in a dialog box.

Electrophysiology controls: Main display controls whether the main displays are plotted or not. Turning off the main displays can be useful during episodic recordings, or when using a slow computer, as it decreases the computational load of the program.

Output controls: Fire output commences the generation of the output sequence selected in Master Setup when that sequence’s generation time is set by the user, not a trigger or acquisition event. ‘Unfired’ is lit and reads ‘Fired’ when an output sequence was successfully generated. ‘Times to run output’ is enabled when the Master Setup Experimental type is set to ‘Acquire immediately and output specified number of times’. Then the output is generated the number of times specified. This option is useful when the same output needs to be run repeatedly, but note that this option is not hardware-timed, so the delay between repetitions is variable. (If precise timing is required, the output desired should be completely specified, however many times it is repeated, in the Master Setup output tab.) The chart shows which outputs are currently set to be generated. The ‘Formatted epochs’ tab shows the outputs in the scale set in the Master Setup output tab whereas the ‘Voltage output’ tab shows the outputs as the raw voltage that will be produced by the DAQ card.

Setup/Start/Stop/Close/Quit Labview: SETUP opens the Master Setup window. START begins the input/output operation, which disables accessibility of certain features in the main window until STOP is pressed. STOP ceases the current input/output operation. CLOSE closes the PackIO program If ‘Quit Labview?’ is checked, pressing the CLOSE button will also quit LabVIEW completely if the full version of LabVIEW and not the runtime is being used.

### PackIO: Master Setup - Analog Input tab

The Master Setup window (**Fig. 2**) controls which channels are recorded, what outputs are generated, and how digital lines are set when the START and STOP buttons are pressed in the main window. It also controls how and which channels are displayed on each of the seven main displays. This window automatically opens when PackIO is initially run. While this window is open, no other LabVIEW windows can be accessed.

**Figure 2:**
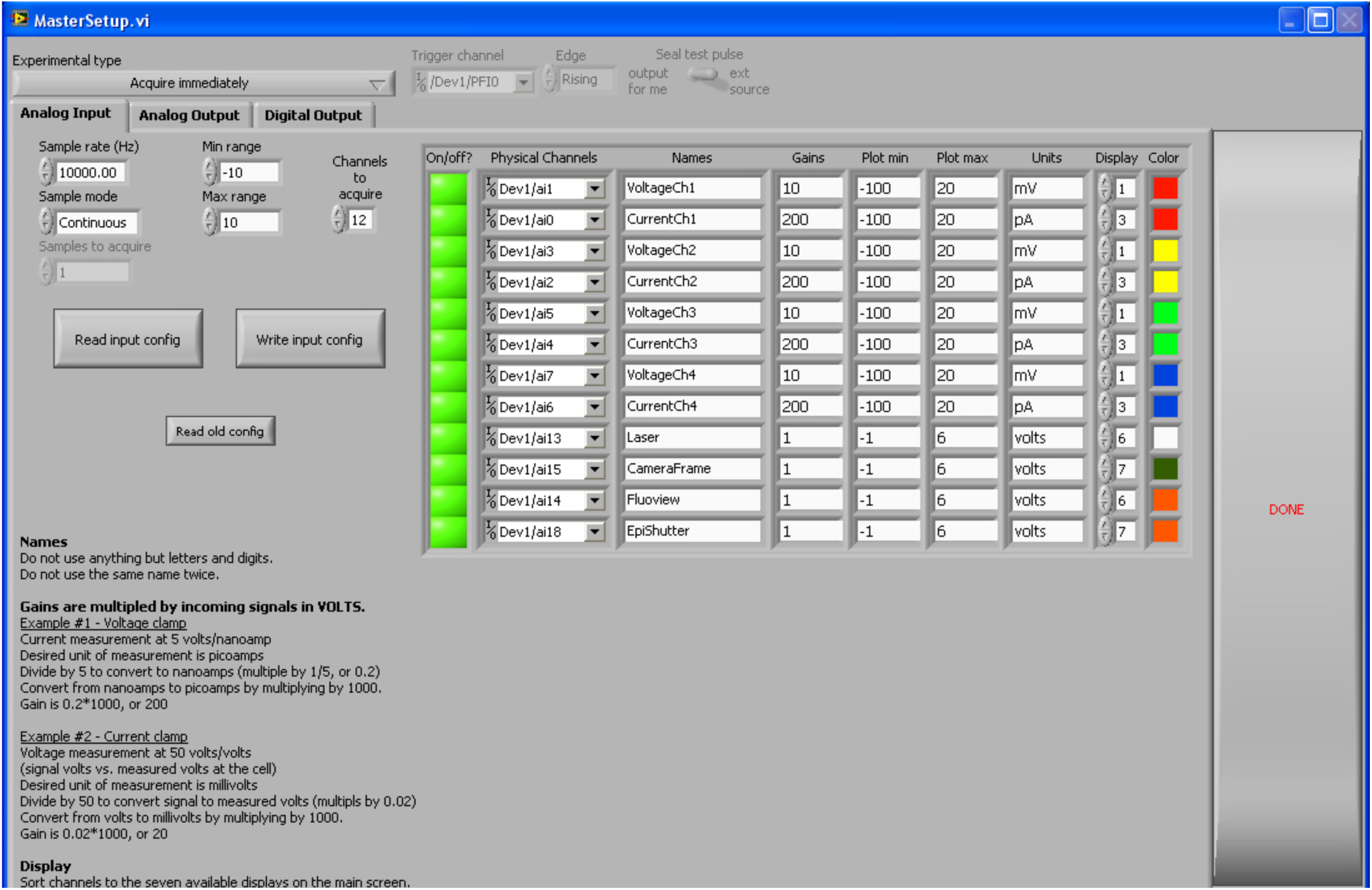
Master Setup analog input window of PackIO. Analog acquisition protocols are designed here.

At the top of the Master Setup window is a critical dropdown box labeled ‘Experimental type’ which controls what type of acquisition, generation, or both is performed as described below:

- ‘Acquire immediately’: Acquire the specified channels when START is pressed.
- ‘Acquire upon external trigger’: Acquire the specified channels after START is pressed and when the trigger condition, as specified by the controls to the right, are met.
- ‘Start acquisition and output immediately’: Acquire the specified channels and generate the specified outputs when START is pressed.
- ‘Synchronize acquisition and output on external trigger’: Acquire the specified channels and generate the specified outputs after START is pressed and when the trigger condition, as specified by the controls to the right, are met.
- ‘Acquire immediately and output upon user input’: Acquire the specified channels when START is pressed. Generate the specified outputs when the user presses the ‘Fire output’ button in the main window.
- ‘Acquire immediately and output upon external trigger’: Acquire the specified channels when START is pressed. Generate the specified outputs when the trigger condition, as specified by the controls to the right, are met.
- ‘Acquire immediately and output specified number of times’: Acquire the specified channels and generate the specified outputs the number of times specified in the main window output section when START is pressed.

Note that acquisition and generation operations which are synchronized between two separate cards, for example between a PCI-6052E and a PCI-6733, require that a RTSI cable be physically installed to connect the two cards inside the computer and appropriately indicated in the NI-MAX configuration software.

The ‘Analog Input’ tab of the Master Setup window controls how many channels are acquired at which frame rate and within what range (upper left.) Those channels can be turned on and off quickly by clicking the green buttons in the column by each channel. The options in this window are self explanatory, except for the gains, which are multiplied by the raw voltage recorded by the DAQ card. Further details and examples are given in the lower left corner of this window. Configurations (left) can be saved and loaded at a later date so that these options need not be specified each time a similar recording configuration is used.

### PackIO: Master Setup - Analog Output tab

The display in the top of the output tab (**Fig. 3**) of the Master Setup window shows which outputs will be generated. It has two tabs which show the output either in the scale set by the Epoch Loader (‘Formatted epochs’, described below), if used, or as raw voltage outputs (‘Voltage output’).

**Figure 3:**
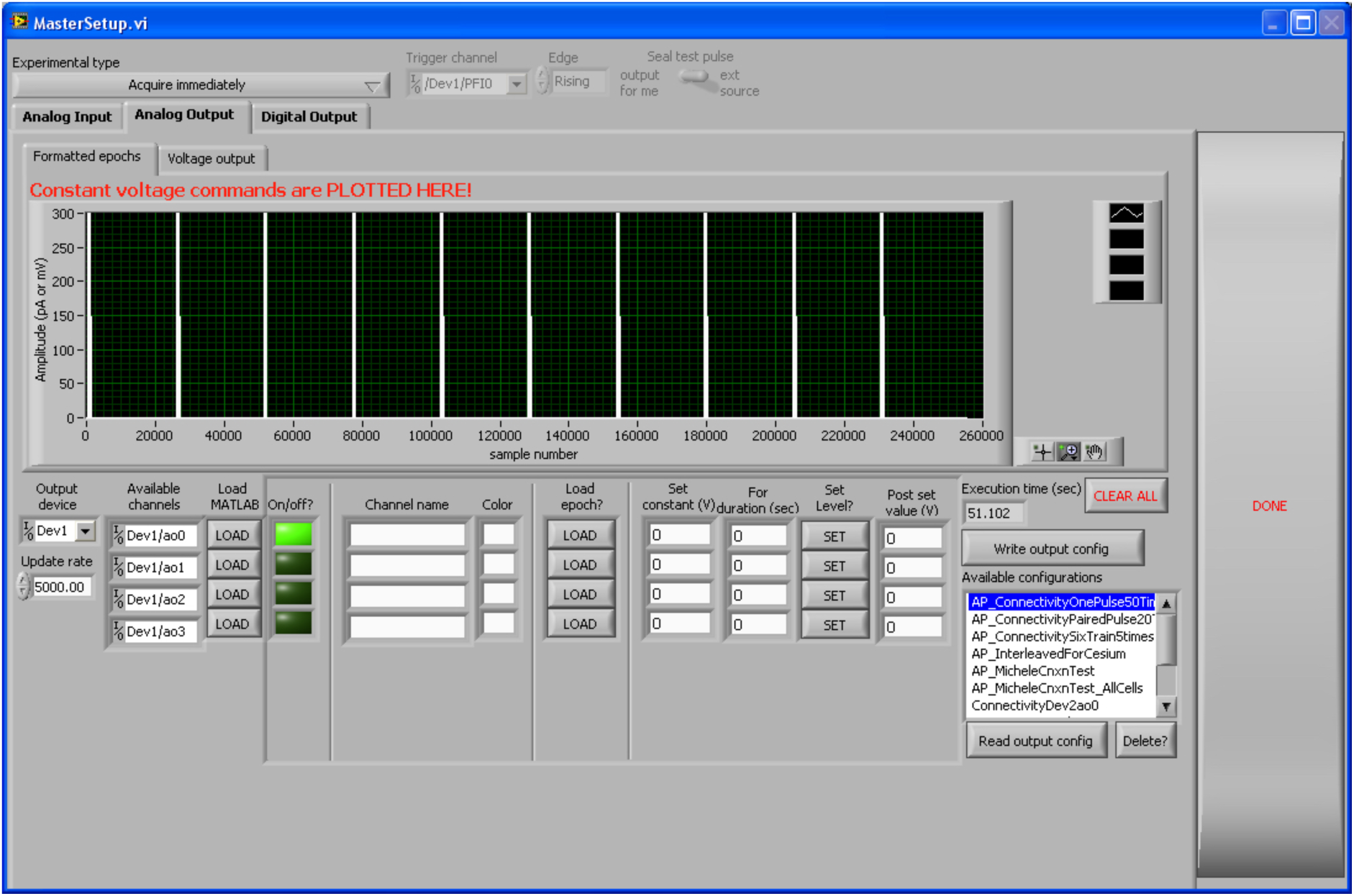
Master Setup analog output window of PackIO. Analog generation protocols are designed here.

Selecting an ‘Output device’ on the lower left will automatically update the available channels. Each channel’s output can be specified by a MATLAB file, an epoch, or a set voltage:

- MATLAB: Use the following commands in MATLAB to write the vector TestRamp to a file TestRamp.dat which can then be loaded into PackIO on a given channel by clicking the LOAD button in the Load MATLAB column: fid=fopen(‘TestRamp.dat‘,‘w’,‘l’);fwrite(fid,TestRamp,‘double’);fclose(fid);
- Epoch: Clicking LOAD in the ‘Load epoch?’ column will open a program (written by Volodymyr Nikolenko), to set various types of stimulation protocols such as steps, ramps, and trains. Gains can be set here so that the user does not have to keep track of voltage to amplitude transformations. Multiple sweeps of a given epoch can also be produced. Individual epochs can be saved for later use.
- Set voltage: A constant voltage can be set for a specified amount of time after which that channel will be set to a given ‘post set value’ after the output is generated.
- Configurations can be saved and loaded so that the user does not have to set the appropriate output for each channel every time the same output paradigm is desired.

### PackIO: Master Setup - Digital Output tab

Digital output lines (**Fig. 4**) can be updated when the START and STOP buttons are pressed in the main window of PackIO. These are useful for sending TTL pulses to various pieces of equipment which need to be toggled before or after an acquisition or generation operation. Note that these lines are not hardware timed, so the time delays before the lines are updated are variable.

**Figure 4:**
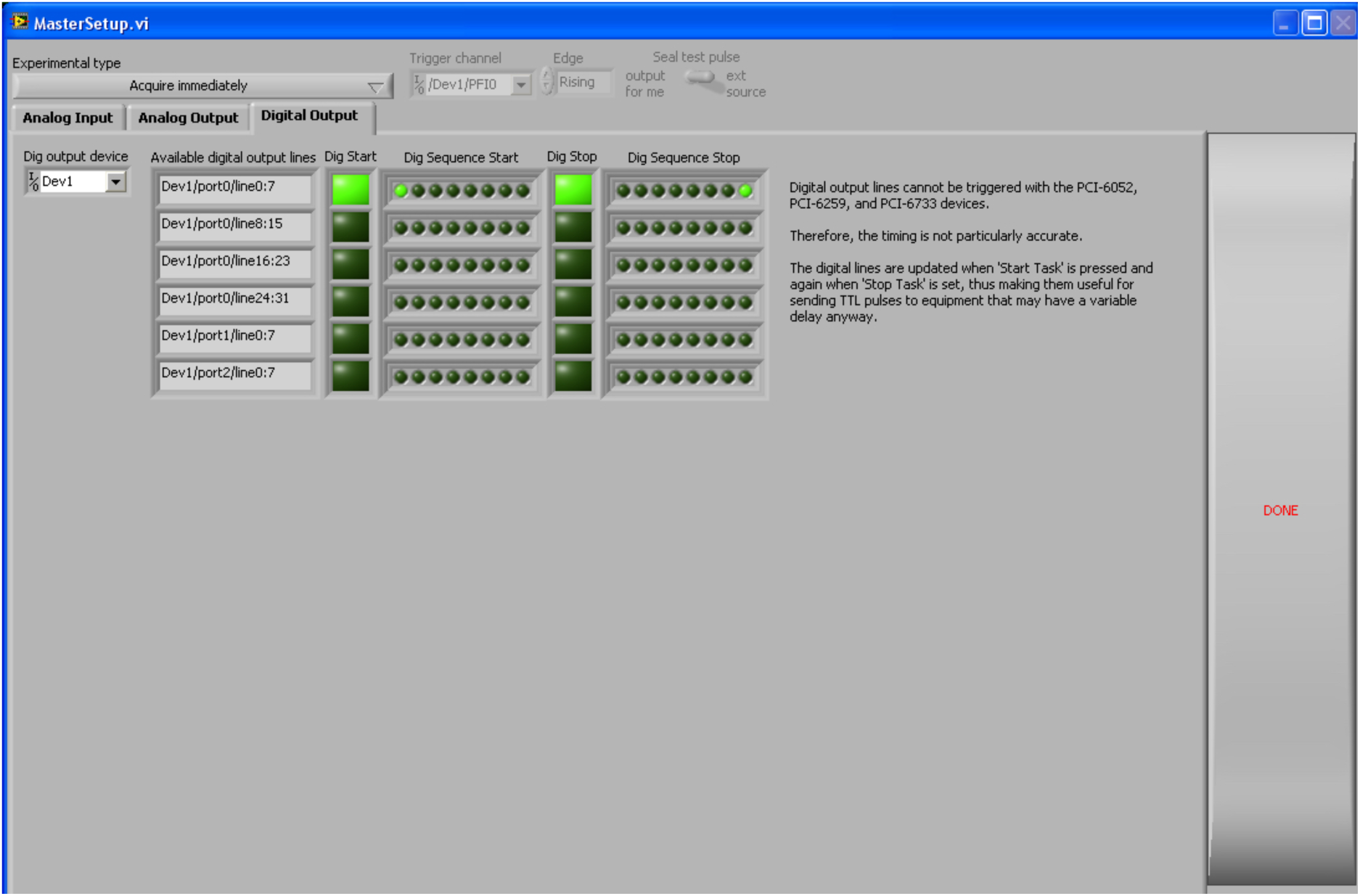
Master Setup digital output window of PackIO. Digital generation protocols are designed here.

### PackIO: Producer/consumer architecture of analog input and output operations

The core of PackIO (**Fig. 5**) that enables the flexible acquisition, generation, and synchronization modes of operation are based on a Producer/Consumer software design pattern^33^. This allows data sharing between multiple loops running at different rates, decouples processes that produce and consume data at different rates, and enables parallelization.

**Figure 5:**
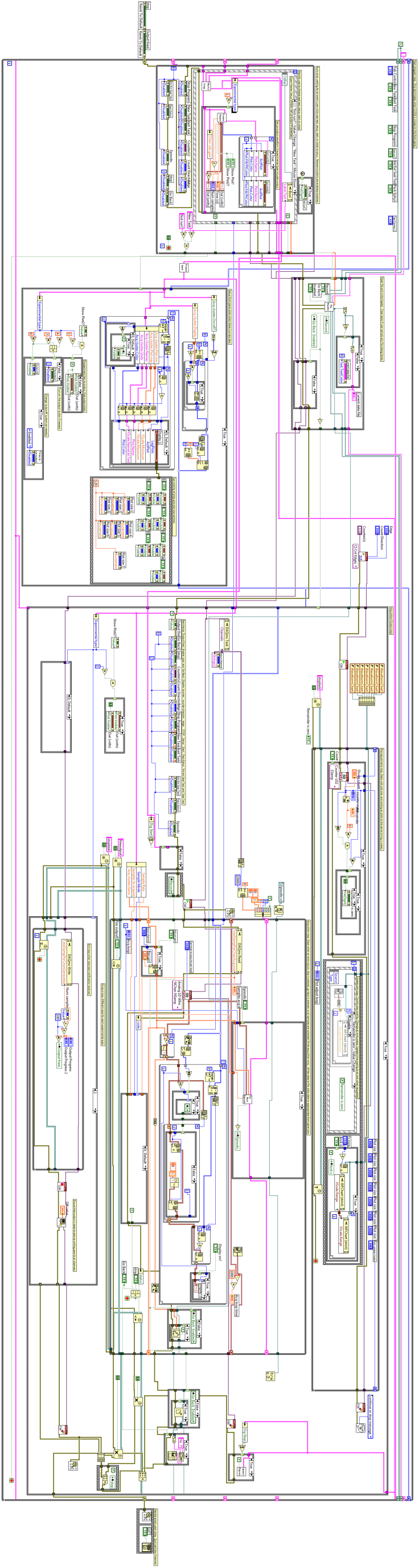
Block diagram (source code) of PackIO’s main acquisition and generation loops. The two large loops on the bottom right of the source code implement the producer/consumer design pattern. This enables the flexibility to switch between different acquisition, generation, and synchronization modes. Code is available online at www.packio.org.

### PackIO: Seal test

The seal test window (**Fig. 6**) allows the user to test the access and input resistance of a recorded cell by calculating the response current to square pulse voltage commands. It can be operated in two modes: ‘Internal’ or ‘External’. Internal mode, selected by clicking the ‘Internal Seal Test’ button near the top left of the window, produces pulses of the specified parameters to the selected ‘Output Channel’. This should be connected to the command of the electrophysiological amplifier to drive the cell. If internal mode is not selected, external mode (so called because the seal test pulses are provided by the amplifier) is used. The sync channel from the electrophysiological amplifier should be connected to the selected ‘Sync Channel (external output)’. The voltage and current channels for the recorded cell need to be appropriately set with the correct gains. The number of samples acquired per run loop of the seal test subroutine can be set, as well as the sample clock rate at which to acquire samples. The pulses per acquisition and actual clock rate are then displayed above the graph on the right. The seal resistance is displayed by the needle and the large grey box directly to the left of the START and STOP buttons (which commence and cease acquisition and, if in internal mode, generation of the seal test pulses). The displayed access and input resistances are correct after the appropriate recording configuration is obtained. The graph on the right shows the response to the seal test pulses. The voltage pulses can also be displayed by toggling the ‘Voltage On/Off?’ button on the upper left of the graph. Configurations can be saved for later use and loaded by selection from the dropdown box.

**Figure 6:**
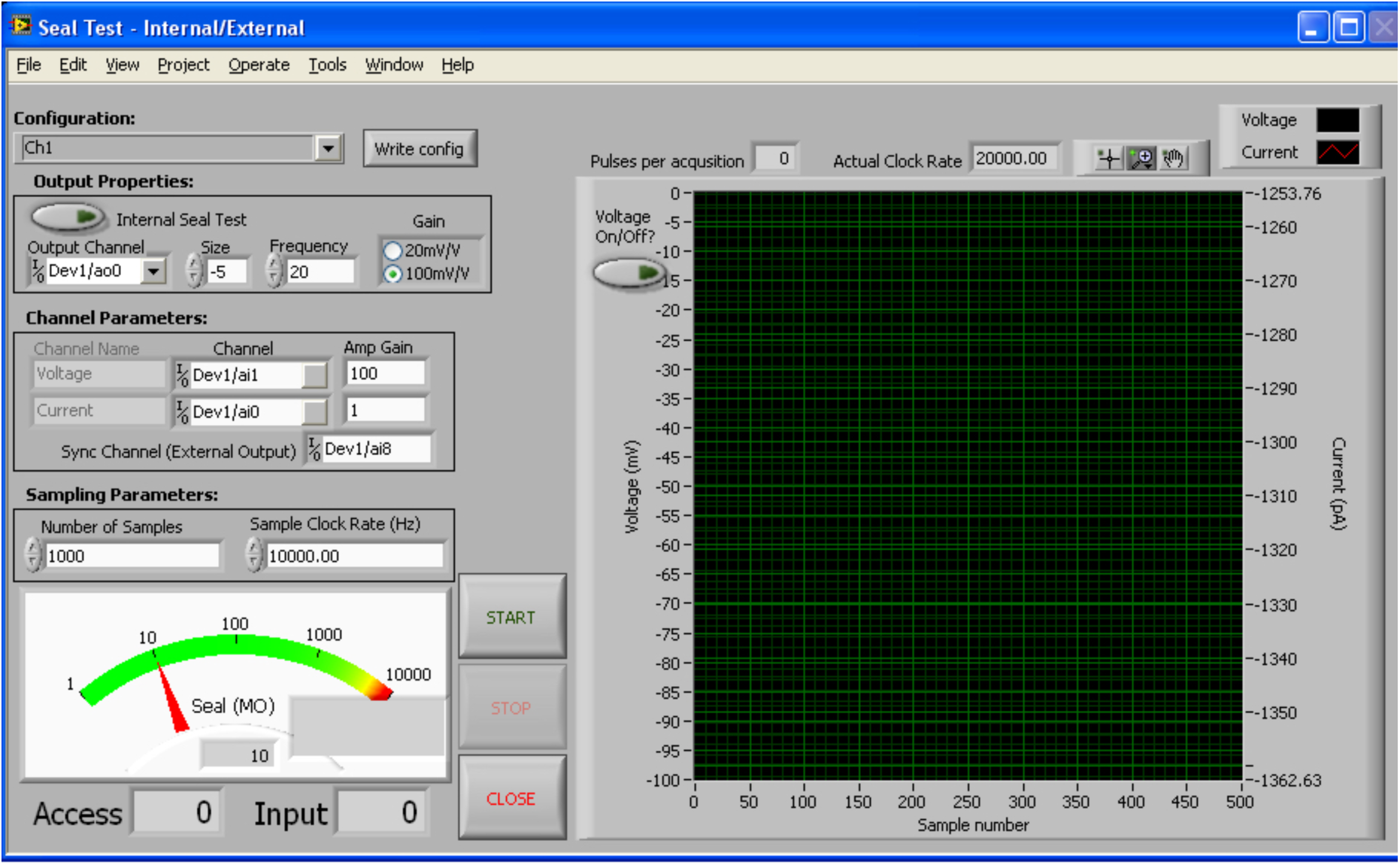
Seal Test. This module enables testing the resistance of a pipette and/or neuron during electrophysiological recordings.

### PackIO: PackEphys

PackEphys (**Fig. 7**) is a electrophysiology recording and control program that can automatically probe the electrophysiological characteristics of cells. After setting the correct channels and gains, the user should select the ‘Threshold’ protocol and a low initial value for the scale factor and then click START. After prompting for a file name, the program will output a square pulse of the scaled amplitude. The user should increase the scale until the recorded cell produces at least one action potential. Then the user can select other protocols to apply to the cell at the same scale. A file produced by ‘All parameters’ can be automatically analyzed by PackEphys.m, a MATLAB script available from the <u>Yuste lab website</u>. The electrophysiological characteristics that can be obtained are based on those from reference 34.

**Figure 7:**
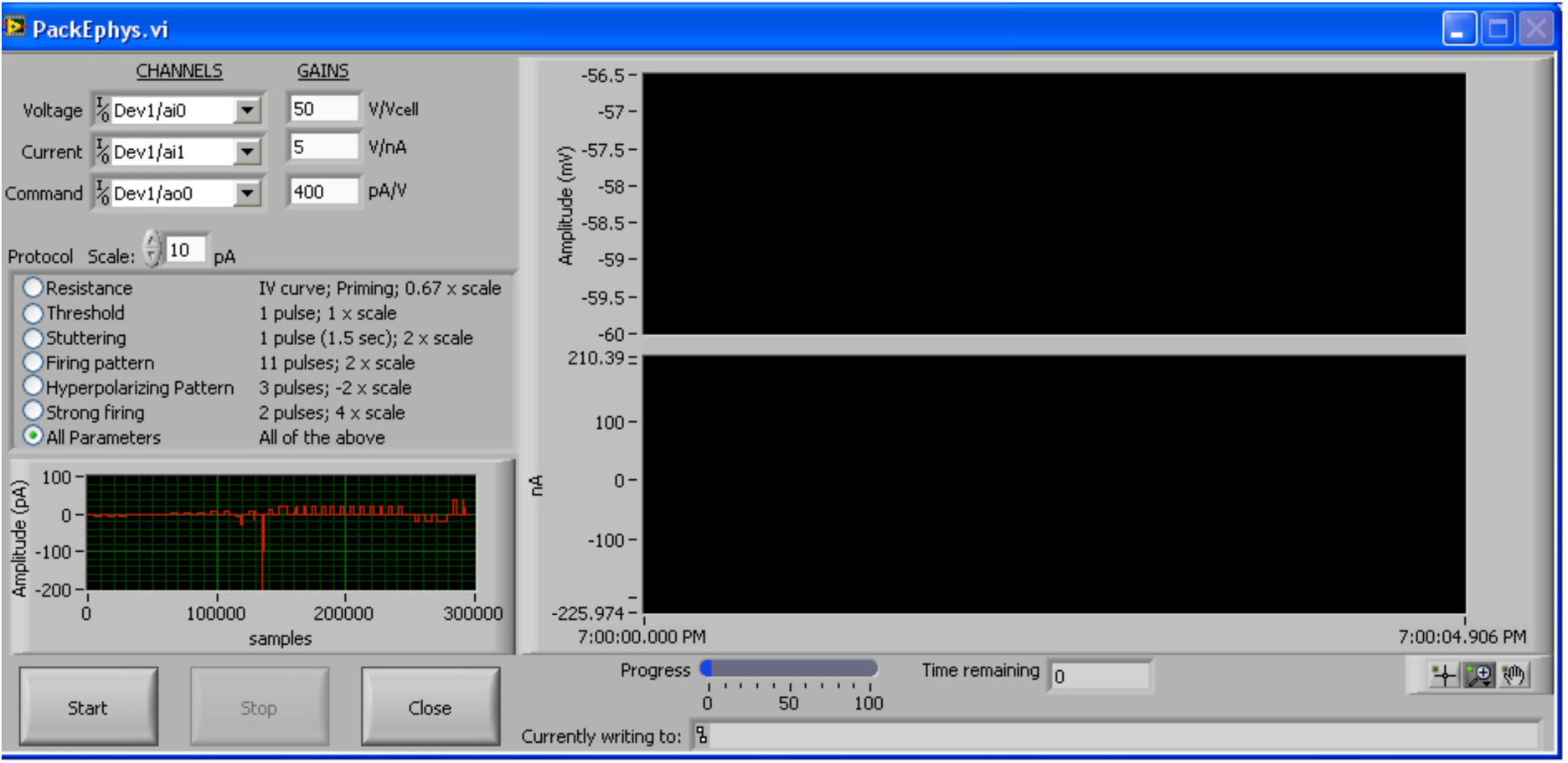
PackEphys. This module enables electrophysiological characterization of neurons.

### PackIO: Episodic acquisition mode

Data being acquired can be displayed in an episodic fashion by clicking the ‘Episodic display’ button on the main window of PackIO to bring up the Episodic window (**Fig. 8**). Available channels will be automatically listed based on the channels that have been set for acquisition in the Master Setup window. Channels the user wishes to view in an episodic fashion should be selected by clicking the episodic button to the left of the name of that channel. An average can be accumulated across sweeps for a given channel by clicking the appropriate button in the ‘Average?’ column. Individual sweeps are displayed in the two graphs at the top right while accumulated averages are displayed in the two graphs at the lower right.

**Figure 8:**
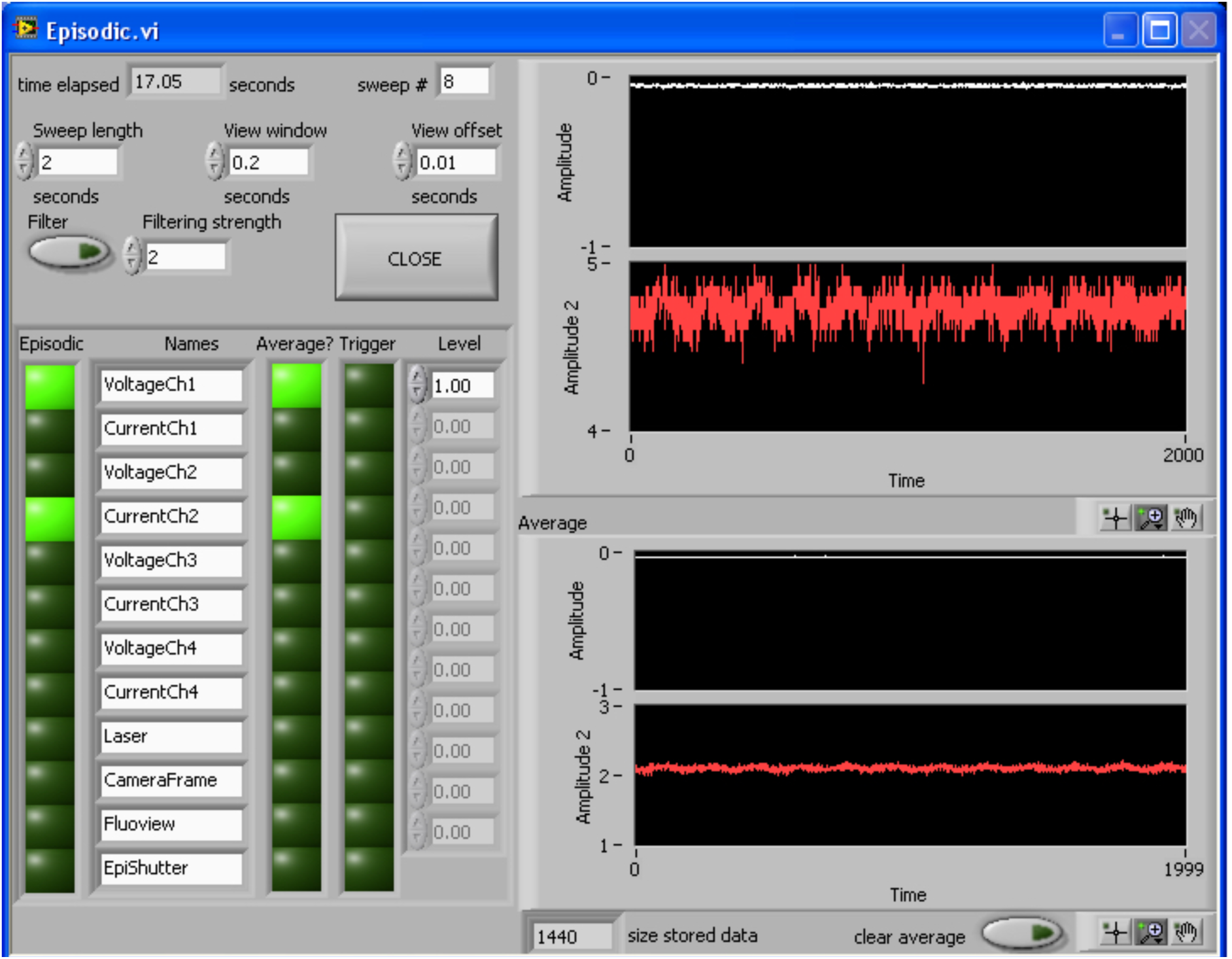
Episodic. This window display data in a trial-wise, or sweep-by-sweep, fashion.

Episodes can be defined either by a set window or a trigger. If no channels are selected as a trigger by clicking one of the buttons in the ‘Trigger’ column, the episodes will default to occur at the intervals specified by the ‘sweep length’ at the top left of the window. If a trigger is selected, when the trigger level is met on the selected trigger, an episode will start. The length of an episode is set by the ‘sweep length’ (in seconds) and the length of the data displayed is set by the ‘view window’ (in seconds). A ‘View offset’ (in seconds) can be set to control the time the view window begins within a given episodic sweep. The total time elapsed and episodic sweep number are displayed at the top of the window.

### EphysViewer

In order to view and analyze data produced by the system, we have created EphysViewer (**Fig. 9**), a graphical interface in MATLAB that has both general features for reading and displaying any binary file and also has features allowing for optimized viewing of data generated by PackIO and PackEphys.

**Figure 9:**
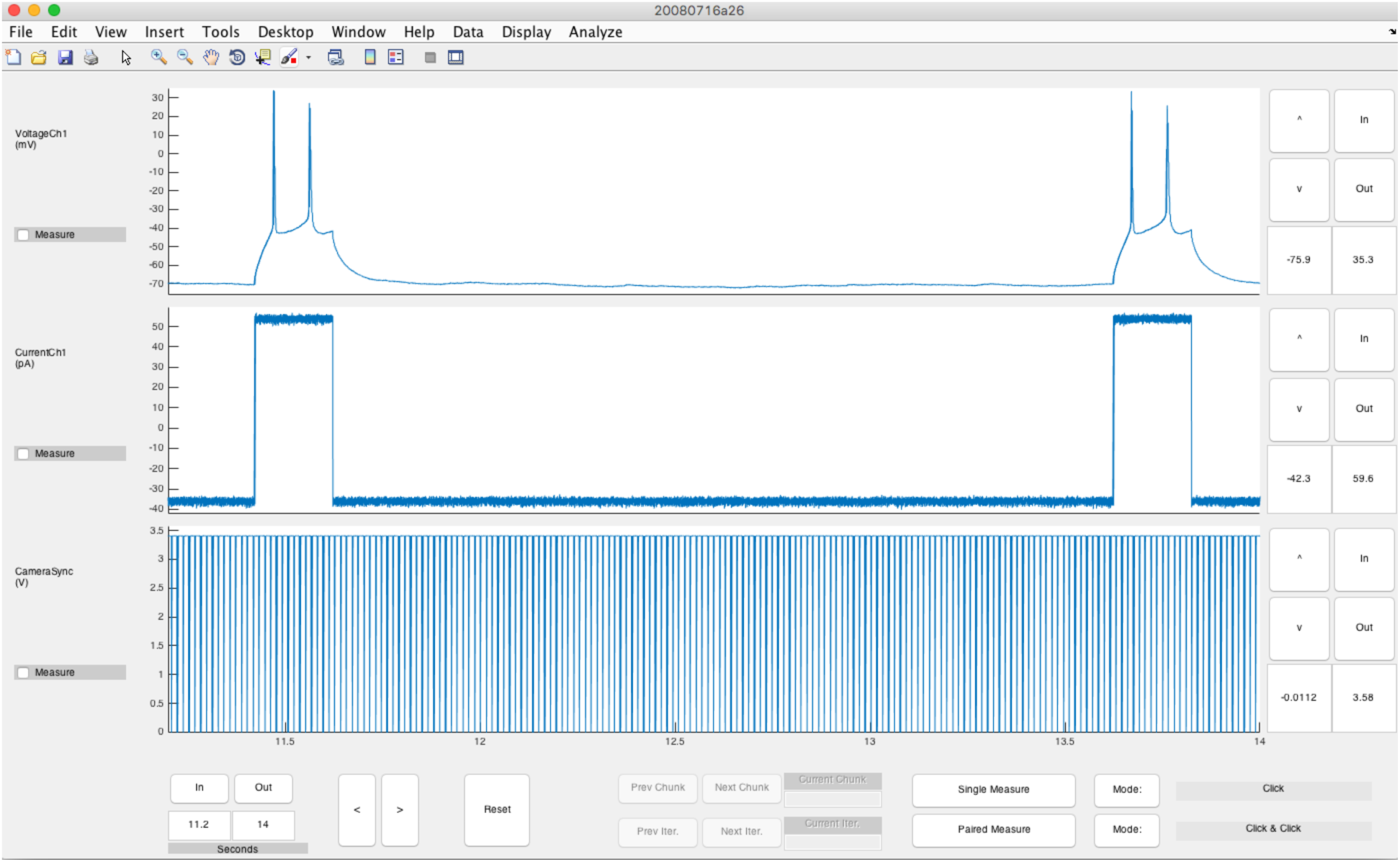
EphysViewer. This GUI allows visualisation and analysis of data acquired with PackIO. In this example, the imaging frame times from a camera and *in vitro* whole-cell patch-clamp electrophysiological data are recorded simultaneously.

### EphysViewer: Display

EphysViewer is optimized to zoom and pan traces more efficiently than in native MATLAB display figures, and for displaying multiple synchronized channels simultaneously. This is achieved by displaying downsampled versions of the data depending on the level of granularity that can actually be visualised. The total number of points displayed is always within an order of magnitude of a visually optimized value regardless of the zoom level. This drastically reduces the time required to draw the data on the screen relative to native MATLAB displaying and zooming. The downsampling is performed immediately after loading the data file from disk prior to display. The decimation method is to save a maximum and minimum value for each decimated set of time points, rather than simply a mean of that span, which led to improved data visualization. A collection of such max-min decimations at different orders of magnitude is stored in the GUI data to access dynamically as zooming or panning is requested. The end result is a display that is visually indistinguishable from native MATLAB plotting but is faster and less computationally intensive to render.

EphysViewer can be used to view and analyze an arbitrary number of simultaneously recorded data channels, though practically greater than 10 channels may be difficult to display. The viewer labels each channel with a text label and maintains that label throughout viewer utilization. Each channel has individually controllable amplitude/y-axis zoom but range changes to the time/x-axis are always synchronized across all channels so that multi-channel data is always viewed in synchronized format. Zooming can be done with mouse clicks or with text entry of start-stop points.

### EphysViewer: File formats

EphysViewer loads files in two formats: *.daq files created by the MATLAB data acquisition toolbox and *.paq 32-bit binary files created by PackIO.

### EphysViewer: General-Purpose Analysis of Continuous Traces

EphysViewer allows for logging and export of measurements of values of the displayed traces at user-specified points. The user may decide to record values at certain points by clicking on them and may choose to record the x and y values of anywhere between one and all channels at that x point. Furthermore, paired points may be recorded and compared so that x and y values and x and y differences may all be obtained. Any measure, either single point-based or two-point based, will be displayed in independent display windows in table format already labeled by channel label and channel number. The user may enter comments as well. At any point, the accumulated measures may be exported to.mat or.xls files or may be exported to currently open MATLAB or Excel sessions.

If more customized or specialized analyses are desired, the user may export the full resolution version of the currently visible data to the MATLAB session. This may include the entire recording if the viewer is simply reset to full zoom out. Visible data may also be exported to disk in MATLAB.mat format.

### EphysViewer: Detection and averaging

EphysViewer can detect peaks or dips (troughs). The user specifies the channel in which to perform the detection, the amplitude of the events and the maximum and minimum durations of events as well as the time window in which to detect or ignore peaks/dips and then a second channel to synchronize to these events to allow for measurement. A new window is then created showing overlaid triggering events, averaged triggering events, overlaid events from the second channel and averages and zeroed averages from that channel. These events are then exportable to the current MATLAB base workspace.

This can be used to make various event-triggered averages including action potential-triggered measures of post-synaptic cell traces to search for synapses in paired recordings. Alternatively, a channel recording machine-driven stimuli, such as those from PackIO can be used as the trigger and cellular voltage or current responses can be overlaid and averaged.

Specific template-matching is available for EPSPs, EPSCs and cell-attached action potentials is available. A correlation-based method is used to search for matches in the recorded data to previously-recorded natural versions of each of these event types. Again, these events can be exported for further analysis. Other templates can be added to the library by users with moderate MATLAB programming skill.

### EphysViewer: Imaging and Stimulation-Specific Features

A number of features are built into the program which are specifically helpful for simultaneous calcium imaging and stimulation experiments as described above. First, the datastream can be subdivided into “chunks” (or trials) based on user-specified signal dynamics within the data itself. As an example we use voltage changes in the stimulation channels to specifically view one full iteration at a time of the stimulation paradigm used. This functionality is extremely useful for efficiently reviewing findings from a full recording/stimulation session. Other types of “chunks” can be determined using any criteria the user desires, assuming they are able to be detected from value changes in data channels.

EphysViewer can also interact with imaging data. Images can be loaded and displayed and contours of cell outlines can be overlaid. Target sequences can be additionally overlaid while viewing the stimuli and cellular responses at once. A full experiment can then thereby be visualized, especially in chunks.

### Python compatibility

The paq2py (https://github.com/llerussell/paq2py) module imports data acquired with PackIO into Python.

## Discussion

At the time this software was initially developed (2007), no software was freely available that could easily and flexibly synchronize electrophysiological recordings with other pieces of equipment (cameras, lasers, galvanometers, etc.). The goal of PackIO was to develop a comprehensive platform capable of performing any data acquisition or generation operation that could be completed with National Instruments hardware in use at the time. PackIO has grown into a system that is useful for any experiment requiring acquisition and generation of data and can be triggered in several modes.

## Methods

### Website

www.packio.org links to all open-source code and necessary software packages.

### Installation

1. Download and install (<u>32-bit</u> or <u>64-bit</u>) LabVIEW Runtime Engine (free).
2. Download and install <u>DAQmx drivers</u> (free).
3. Download and run <u>32-bit</u> or <u>64-bit</u> PackIO.

A full installation of LabVIEW is required to edit the source code. Many universities have a site license for this programming environment.

### Supported hardware

Most National Instruments DAQ cards should work. The following cards are known to work:

- PCIe-6343
- PCI-6259
- PCI-6052e
- PCI-6711
- PCI-6713
- PCI-6733
- USB-6009
- USB-6211
- USB-6343
- USB-6251

### Software license

PackIO and EphysViewer are released under the GNU General Public License.

## Acknowledgments

We thank the beta testers of the software from our laboratories. We also thank Volodymyr Nikolenko, who contributed the epoch loader module; Mor Dar, who contributed some features to seal test; and Carmen F. Fisac, who contributed the auto-incrementer available in version 273. This work was supported by the NEI (DP1EY024503, R01EY011787), NIMH (R01MH101218, R01 MH100561), DARPA SIMPLEX N66001-15-C-4032, the Wellcome Trust, the Gatsby Charitable Foundation, the European Commission (Marie Curie International Incoming Fellowship grant no. 328048), the European Molecular Biology Organization, the Medical Research Council and the European Research Council.

### Author Contributions

A.M.P. authored PackIO. B.O.W. authored EphysViewer with some contributions by A.M.P. A.M.P. and B.O.W. wrote the manuscript with input from all authors. R.Y. and A.M.P. supervised the project.

